# Alpha-, beta-, and gamma-diversity of bacteria varies across global habitats

**DOI:** 10.1101/2020.05.15.097758

**Authors:** Kendra E. Walters, Jennifer B.H. Martiny

## Abstract

Bacteria are essential parts of ecosystems and are the most abundant organisms on the planet. Yet, we still do not know which habitats support the highest diversity of bacteria across multiple scales. We analyzed alpha-, beta-, and gamma-diversity of bacterial assemblages using 11,680 samples compiled by the Earth Microbiome Project. We found that soils contained the highest bacterial richness within a single sample (alpha-diversity), but sediment assemblages were the most diverse at a global scale (gamma-diversity). Sediment, biofilms/mats, and inland water exhibited the most variation in community composition among geographic locations (beta-diversity). Within soils, agricultural lands, hot deserts, grasslands, and shrublands contained the highest richness, while forests, cold deserts, and tundra biomes consistently harbored fewer bacterial species. Surprisingly, agricultural soils encompassed similar levels of beta-diversity as other soil biomes. These patterns were robust to the alpha- and beta-diversity metrics used and the taxonomic binning approach. Overall, the results support the idea that spatial environmental heterogeneity is an important driver of bacterial diversity.

## Introduction

Bacteria are the most abundant organisms on the planet [1]. The richness and composition of this diversity influences ecosystem functioning, whether in host-associated communities, soils, or oceans [2–7]. Nevertheless, we have yet to answer a number of basic questions about bacterial diversity, including “Where does the highest diversity of bacteria reside on the planet?” More broadly, evaluating geographic patterns in biodiversity across habitats and spatial scales can illuminate the processes influencing and consequences of biodiversity [8–10].

While many studies document spatial patterns of bacterial diversity, most are restricted to a particular geographic region or habitat, such as soil, sediment, or water [11–13]. To understand global trends, however, studies that analyze diversity across habitats and geographic regions are needed. Combining data from independent projects is oftentimes infeasible because community variation can be caused simply by differences in methodology. The Earth Microbiome Project (EMP) is a global dataset of bacterial diversity comprising 27,751 samples from 97 studies from a wide range of habitats and geographic regions that are processed in the exact same way [14]. This database thus provides an opportunity for a rigorous comparison of bacterial diversity across the globe.

A recent overview from the EMP noted, as has previously been observed, that communities of free-living bacteria are more diverse than host-associated bacteria [15–17]. For example, soil and sediment samples were found to have higher alpha-diversity than animal gut or skin microbiomes. We expand on this initial alpha-diversity analysis by additionally evaluating beta- and gamma-diversity across spatial scales while mitigating for unevenly spaced samples (S1 Fig.). We also take the opportunity to use the large size of the EMP dataset to test whether different diversity metrics and taxa definitions influence our understanding of microbial diversity patterns.

We specifically ask: which habitats across the globe support the highest levels of bacterial diversity? We consider three inter-related aspects of biodiversity: alpha-, beta-, and gamma-diversity. We measure alpha-diversity as the observed richness (number of taxa) or evenness (the relative abundances of those taxa) of an average sample within a habitat type. We quantify beta-diversity as the variability in community composition (the identity of taxa observed) among samples within a habitat across the globe [18]. Finally, we calculate gamma-diversity as the total observed richness in a habitat across the globe.

We test several predictions about relative, not absolute, diversity patterns because, even in this large dataset, bacterial diversity remains undersampled. First, we predict that sediment and soil support the highest alpha-diversity within a single sample. These habitats are known to have relatively high bacterial diversity, although their relative rankings have not yet reached a consensus [14–17]. Second, we expect that soil, sediment, inland water, and biofilm/mat habitats will exhibit high beta-diversity. These habitats are spatially separated with less dispersal or mixing than air or marine water. Finally, we predict that soils and sediments will exhibit high global, or gamma-, diversity as they are expected to have both high alpha- and beta-diversity.

Within the soil habitat, we hypothesize that soils from biomes higher in plant diversity and productivity (e.g., forests and grasslands) support higher alpha-diversity than soils from biomes with low diversity and productivity (e.g. tundra and deserts) [19–21]. Of course, these biomes do not directly influence diversity, but they are defined based on abiotic factors [22], such as temperature or precipitation, that do influence diversity. Further, we expect that agricultural soils will exhibit the lower beta-diversity than other biomes as common practices (pesticides, tilling, and fertilizer use) and the low diversity of crop plants influences community composition [23,24]. We also compare the relationship between diversity and biomes to those between diversity and pH or temperature to assess whether plant diversity or abiotic conditions are more influential on bacterial diversity. Our results show that the habitat that is the most diverse depends on spatial scale (alpha-, beta-, or gamma-diversity).

## Materials and Methods

Bacterial 16S (V4 region) sequence data and associated metadata were downloaded from the Earth Microbiome Project (EMP) on September 1, 2016. Sample processing, sequencing, and core amplicon data analysis were performed by the Earth Microbiome Project (www.earthmicrobiome.org), and all amplicon sequence data and metadata have been made public through the data portal (qiita.microbio.me/emp). Data available from: https://doi.org/10.1038/nature24621 [14]. For most of the analyses, we used the EMP closed-reference (Greengenes 13.8) OTU database classified at 97% sequence similarity to reduce computational time. The database contains 27,751 samples, with a median depth of 54,091 sequences per sample. We excluded archaea from the analysis because, relative to bacteria, they make up a small portion of any given community.

### Habitat designations

We used the EMP Ontogeny metadata to classify the habitat and, for soil, biome of each sample based on the EMP metadata (Fig. 1). When the existing metadata were unclear, we used the latitude and longitude coordinates to assess the environmental context. Samples with insufficient data about their location or habitat were removed from the analysis. We only retained samples that could be classified into one of the following habitats: soil, sediment, marine water, inland water (e.g., rivers and lakes), air, and biofilms/mats. These habitat types were chosen because they represent a wide range of environmental conditions and are well sampled within the EMP dataset. Within the soil habitat, we further classified samples into forest, hot desert, cold desert, grassland, shrubland, tundra, and agricultural soil. We also classified inland water and sediment samples as saline and non-saline. See supplemental materials for descriptions of sample locations (S1 Appendix).

**Fig 1.**
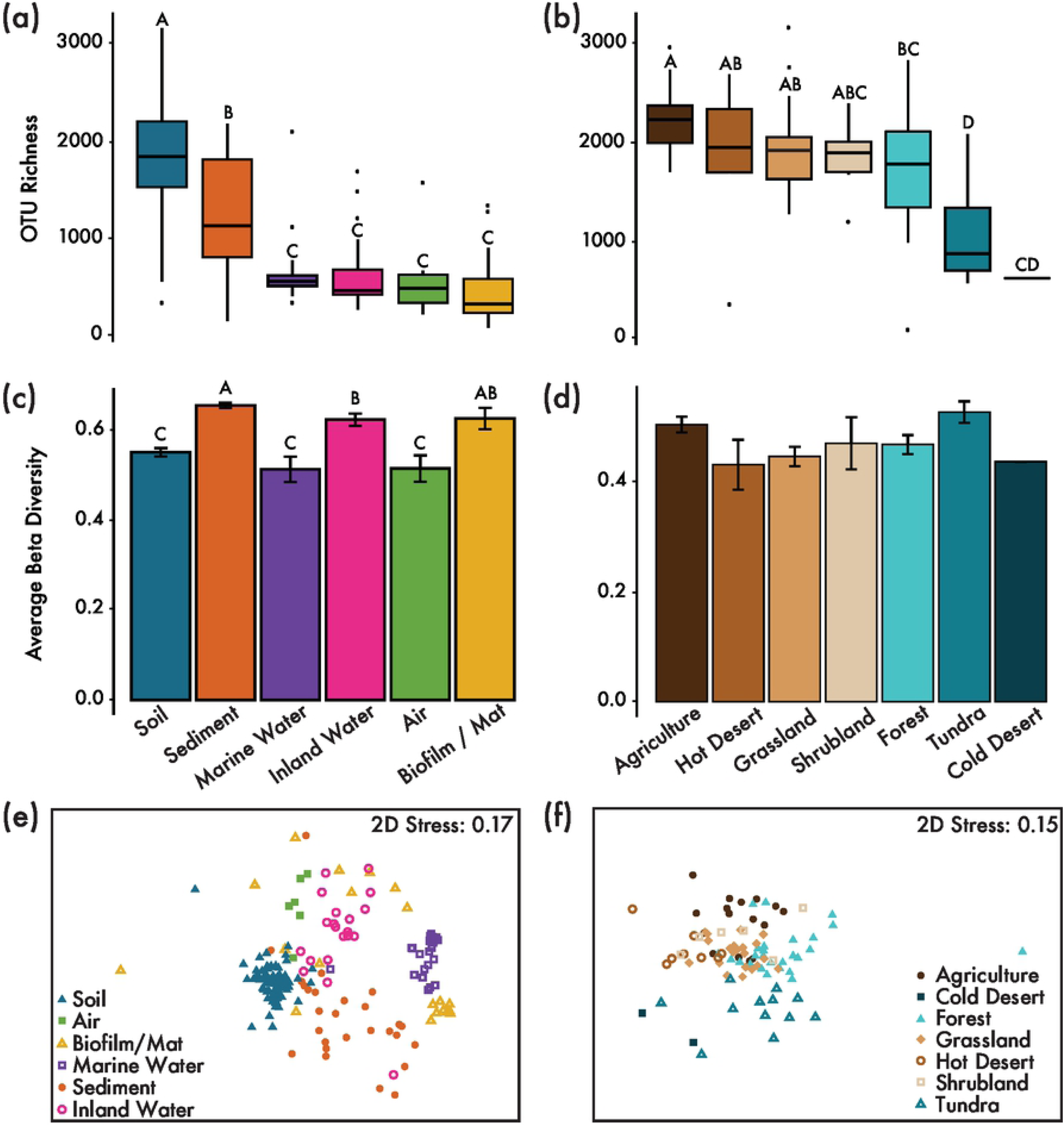
Alpha- and beta-diversity patterns. Alpha- and beta-diversity per habitat for all geoclusters used in study (A, C, and E) and per biome for soil geoclusters (B, D, and F). (A and B) Boxplot of alpha-diversity (OTU richness). (C and D) Mean beta-diversity (distance from centroid) ± standard deviations. (E and F) NMDS of geoclusters.

### Alpha-diversity analysis

To account for differences in sequencing depth, the samples were rarefied to 15,000 sequences with 1,000 resamplings in QIIME [25]. All samples with less than 15,000 sequences were removed leaving a total of 11,680 free-living (non-host system) samples. For each resampling, we calculated 24 alpha-diversity metrics on the rarified OTU table in QIIME (Fig. 2A). These metrics characterized the community in five general ways: observed richness, estimated richness, evenness/dominance, phylogenetic diversity, and coverage of sampling. We used all 24 metrics throughout the alpha-diversity analysis to ensure that our final conclusions were not dependent on the type of metric. We calculated the median value of each metric across the 1,000 replicates.

**Fig 2.**
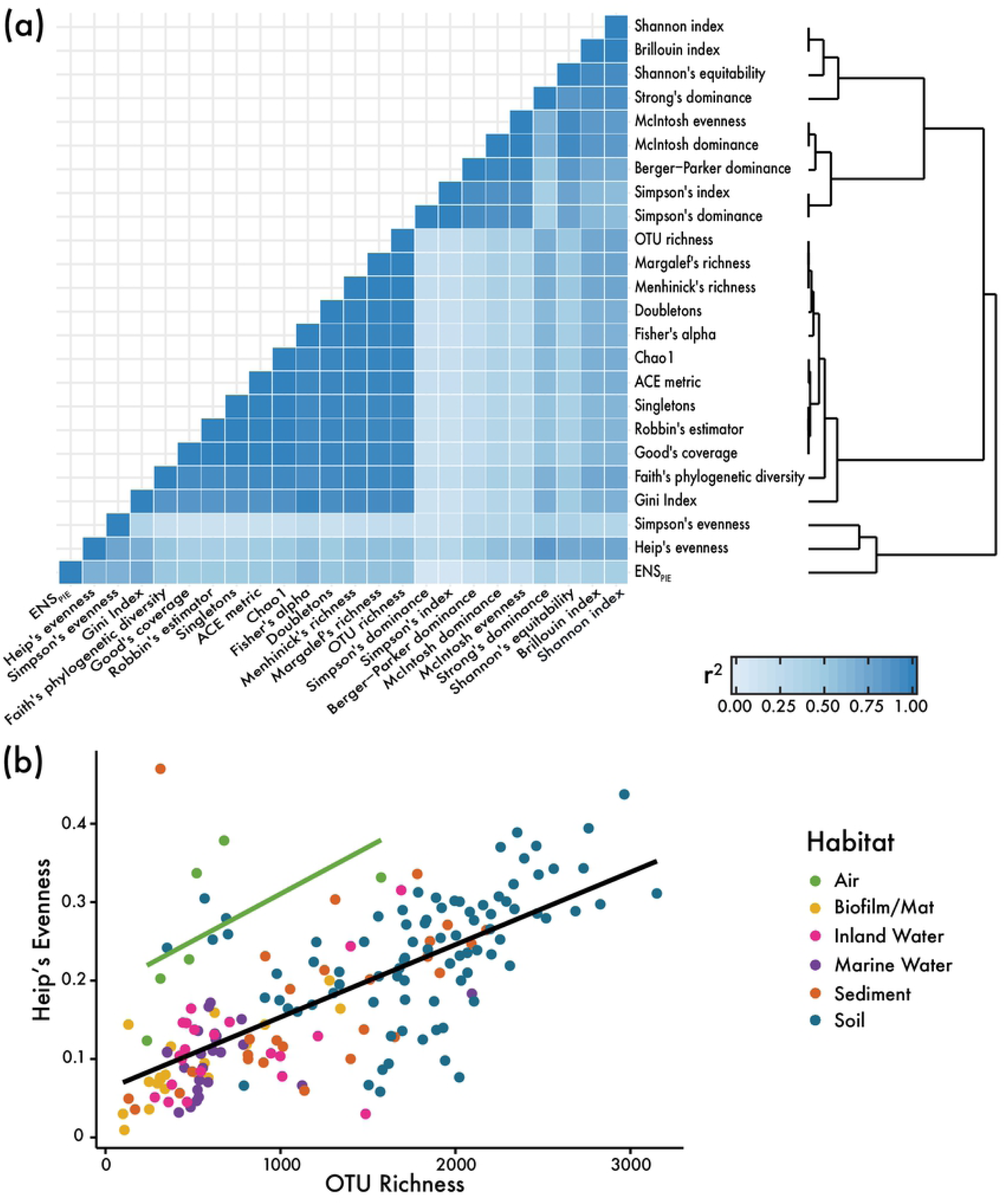
Comparison of diversity metrics. (A) Heatmap shows degree of correlation (r^2^ from linear regression with all EMP samples used in analysis). Dendrogram shows relatedness of metrics based on their correlation strength. Note that the metrics are clustered into two groups: one composed of mainly evenness metrics (top cluster on dendrogram) and one composed of mainly richness metrics (bottom cluster on dendrogram). Simpson’s evenness, Heip’s evenness, and ENS_pie_ fall outside of those two clusters. (B) Dot plot showing relationship between Heip’s evenness and OTU richness metrics for geoclusters of each habitat. The green line is a linear regression for air geoclusters, and black line is a linear regression for all geoclusters except for air. Excluding air samples, habitat is not a predictor of evenness (ANCOVA: p-values < 0.00001 [OTU richness], 0.0823 [habitat]). When air samples are included, habitat is a significant predictor of evenness (ANCOVA: p-values < 0.0001 [OTU richness, habitat]).

To minimize the effect of unevenly spaced samples, we averaged the alpha-diversity of the samples within a single geocluster. Many of the samples are highly clumped such that some geographic regions contribute unequally to the habitat’s diversity. Geoclusters (n= 172) were formed by clustering samples of the same habitat type within 110 km of each other using hclust() and cutree() from package ‘stats’ and rdist.earth() from package ‘fields’ in R [26,27]. We calculated the median of each diversity metric for the samples within each geocluster of the same habitat type. The averaged alpha-diversities were then cube root transformed to achieve normality and homoscedasticity. Finally, we tested for significant differences in alpha-diversity among habitats by performing a one-way ANOVA and Tukey’s HSD in R.

We tested whether the alpha-diversity results depended on the diversity metric used by running a correlation with every pairwise combination of diversity metrics using all 11,680 samples. Likewise, we tested whether our results depended on the resolution of OTU clustering by comparing the 97% similarity OTU table with the single-nucleotide resolution ‘sub-OTUs’ database, Deblur, produced by the EMP. We rarefied the Deblur dataset to 15,000 sequences per samples (1,000 times), calculated Exact Sequence Variance (ESV) richness per sample, and calculated the mean richness across the 1,000 replicates. We then ran a correlation between the OTU richness and ESV richness for every sample present in both datasets (n = 20,474).

To further explore what factors might be driving alpha-diversity, we compared bacterial richness with pH and temperature at each sample site. We used metadata from the EMP, representing pH and temperature taken at the site at the time of sample collection, and from WorldClim, a publicly-available dataset with mean annual temperature averaged from 1970 – 2000 with spatial resolution of 10 minutes. Data available from WorldClim Version2: http://doi.org/10.1002/joc.5086 [28]. We assigned external temperature data to soil samples using latitude and longitude with extract() from package ‘raster’ in R [29]. Temperature and pH were correlated with OTU richness using a second-degree polynomial in R. We tested whether temperature and pH differed among habitats and biomes using an ANOVA in R.

### Beta-diversity analysis

To reduce computation time, we used a subset (150 tables) of the rarefied OTU tables from the alpha-diversity analysis to analyze beta-diversity. The OTU tables were first square root transformed to give more weight to rarer taxa. For each of the 150 rarefied, square root transformed OTU tables, we calculated the median abundance for each taxon across all the samples within a single geocluster. We then calculated a Bray-Curtis dissimilarity matrix for each of the 150 OTU-by-geocluster tables in QIIME. Finally, we calculated the median of the 150 dissimilarity matrices to yield one median Bray-Curtis dissimilarity matrix.

To visualize compositional differences among habitats, we used NMDS in PRIMER6 [30]. We also tested community composition differences among habitats using PERMANOVA in PERMANOVA+ [31]. Because we averaged OTU abundances for all samples of the same habitat type located within the same geographic area (geocluster), beta diversity provides an approximation of the amount of community variation from location to location within one habitat (as opposed to variation from sample to sample within one location).

To compare beta-diversity across habitats, we analyzed the variance within each habitat using the function PERMDISP in PERMANOVA+. To determine if unequal sampling among habitats biased these results, we re-calculated the Bray-Curtis values based on a selection of only 20 geoclusters for each habitat from a rarified OTU-by-geocluster table. We chose 20 geoclusters, because the number included five of the six habitats (excluding air) but avoided the biases expected with sample sizes less than ten [31]. We repeated these subsamplings 100 times and tested for differences in beta-diversity among habitats using betadisper(), the PERMDISP test implemented in the R package vegan [32]. We compared the relative rankings of these rarefied beta-diversity results to the unrarefied results to determine if rarefaction changed the relationships of variance among habitat groups. Specifically, we considered the rarefied results to match the unrarefied results if 95-100 subsampled tests were significant and showed the same beta-diversity rankings (based on mean distance to centroid) as the unrarefied test.

We tested whether the beta-diversity results depended on the diversity metric used by running a correlation with every pairwise combination of nine diversity metrics. For each of the 150 OTU-by-geocluster tables, we calculated nine beta-diversity metrics using vegdist() from package ‘vegan’ in R [32]. We then took the mean matrix (of the 150 matrices) for each diversity metric. We performed a Spearman’s mantel test for every pairwise comparison of the nine averaged beta-diversity matrices using mantel() from package ‘stats’ [26]. To compare Raup-Crick to the other beta-diversity metrics, we calculated the dissimilarity within, but not among, habitats. Raup-Crick is not an appropriate metric when communities do not share the same species pool [33]. To calculate Raup-Crick, we took the mean of the matrices computed with raupcrick(…, chase = TRUE) and raupcrick(…, chase = FALSE) from package ‘vegan’ in R [32] to follow the method recommended by Chase et al. [33]. We calculated one Raup-Crick matrix for each habitat for each of the 150 OTU-by-geocluster tables, took the mean for each habitat across the 150 matrices, and then calculated the mean and SE distance from centroid in PRIMER6 [30]. We used the mean and SE to compare the trends in dissimilarity to those generated with the Bray-Curtis metric.

### Gamma-diversity analysis

To assess gamma-diversity by habitat, we plotted an OTU accumulation curve for each habitat with specaccum() from package ‘vegan’ in R [32] using the 150 OTU-by-geocluster tables. The OTU accumulation curve displays the numbers of geoclusters sampled on the x-axis and observed OTU richness on the y-axis. This plot allowed us to compare cumulative diversity levels across multiple samples distributed across the world. We examined whether habitats likely exhibit different gamma-diversity levels by calculating error bars equal to 1.96 times the standard deviation.

## Results

### Alpha Diversity

Out of the six habitats compared, soils contained the highest observed richness (i.e., number of observed taxa rarefied at 15,000 sequences) for a single sample, with a median of 1,842 taxa (97% OTUs) per sample given this depth of sequencing (one-way ANOVA: P < 0.001; Fig. 1A). Sediments were the second most diverse habitat globally with an average of 1,137 taxa. Marine water, air, inland water, and biofilms/mats had a significantly lower richness (averaging 571, 500, 478, and 342 taxa, respectively) than soils and sediments and could not be distinguished from one another by richness (P < 0.001).

Because past studies indicate that salinity influences bacterial community composition [15], we further tested whether taxon richness varied between non-saline and saline habitats. We found that salinity had no impact on alpha-diversity for sediment (one-way ANOVA: P = 0.516, S2 Fig.) or inland water (P = 0.763, S2 Fig.).

Within the soil habitat, we further compared alpha-diversity among seven biomes (agricultural, grassland, shrubland, forest, hot desert, cold desert, and tundra soil). Within soil samples, richness differed significantly among biomes. Agricultural soils supported the highest richness in a sample, along with hot desert, grassland, and shrubland biomes. Forest soils were significantly less diverse than those biomes, and tundra and cold deserts supported the lowest richness (one-way ANOVA: P < 0.001; Fig. 1B). Notably, the cold desert biome was only represented by two geoclusters (averaging across 117 samples); thus, more data are needed to assess that particular biome’s diversity.

The above results were robust to the alpha-diversity metric used. On a sample by sample basis, 24 alpha-diversity indices, including observed richness, were all correlated with each other (r^2^ = 0.09 – 1.00, P < 0.0001) with a mean r^2^ of 0.63 (Fig. 2A). The metrics grouped into two main clusters. One cluster encompassed the richness/coverage metrics such as OTU richness, Faith’s Phylogenetic Diversity, and Chao1 (r^2^ = 0.86 - 1.00, mean r^2^ = 0.97). The other cluster included the evenness/dominance metrics such as Simpson’s and McIntosh dominance index (r^2^ = 0.31-1.00, mean r^2^ = 0.85). Further, each metric ranked the habitats from highest to lowest alpha-diversity in the same way, with the exception of air. Air communities were more even than other habitats, given their relative richness level (Fig. 2B).

The alpha-diversity patterns were also robust to the taxonomic binning method (Fig. 3). We compared the 97% OTUs and Exact Sequence Variants (ESV) Deblur datasets provided by the EMP. Not only were the metrics strongly correlated (r^2^ = 0.87, P < 0.0001), but, on a sample by sample basis, OTU richness and ESV richness were nearly identical (slope = 1.017, P < 0.0001). However, the relationship between OTU richness and ESV richness varied among habitats (ANOVA: P < 0.0001). In particular, non-saline sediments and inland water demonstrated a higher ESV:OTU richness ratio than other habitats (Fig. 3).

**Figure 3.**
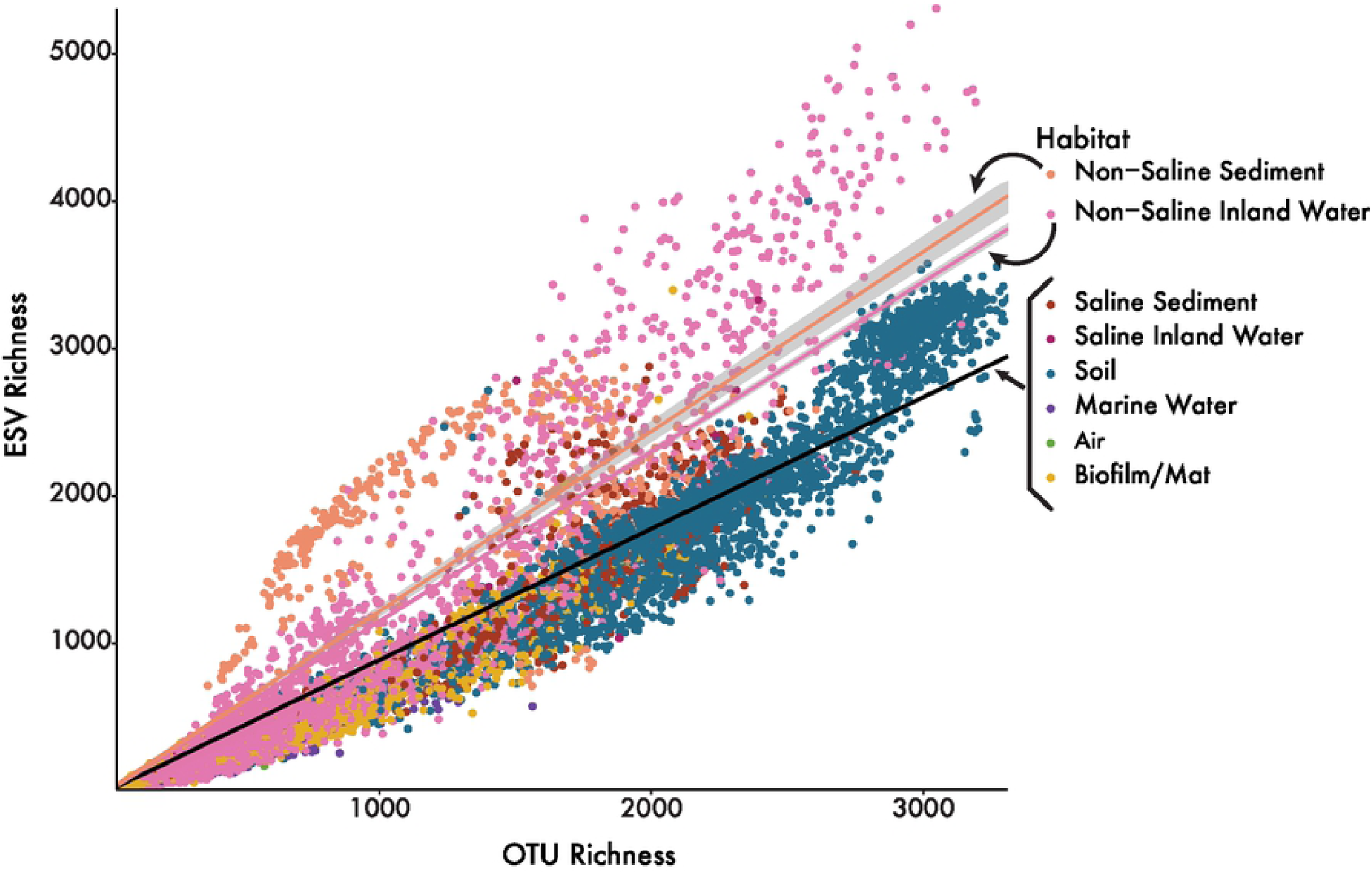
Comparison of OTU and ESV richness. On a sample-by-sample basis, OTU richness and ESV richness are highly correlated (r^2^ = 0.9327, P < 0.0001). On average, for any give sample, ESV richness is equal to 95.26% of OTU richness. However, not all habitats showed the exact same relationship between OTU and ESV richness. Non-saline sediment and inland water samples have significantly higher ESV richness given their relative OTU richness. All samples except for non-saline sediment and inland water have a regression line with a slope of 0.9711 (p-value < 0.0001), shown as the black regression line on the graph. Non-saline sediment samples have a regression line with a slope of 1.220 (P < 0.0001), shown as light orange on the graph. Non-saline inland water samples have a regression line with a slope of 1.152 (P < 0.0001), shown as light pink on the graph.

Taxon richness displayed a weak hump-shaped relationship with pH and a peak in diversity at a neutral pH (P < 0.0001, S3A Fig.). In contrast, taxon richness only weakly correlated with temperature, and this relationship was driven by low-diversity biofilm/mat samples sampled from high temperatures (P < 0.0001, S3C Fig.). Overall, the bacterial alpha-diversity patterns across all habitats were not obviously related to pH or temperature. Most of the samples were, on average, at a neutral pH, with the soil samples more acidic and the biofilm/mat samples more basic (S3B Fig.). Despite this, temperature (ANOVA: P < 0.0001) and pH (P < 0.0001) differed significantly among habitats (S3D Fig.).

Similar to the pattern observed across all habitats, richness within just the soil samples also peaked at a neutral pH (non-linear regression: P = 0.001, S4A Fig.). In contrast, richness in soils also peaked at a temperature around 10°C (P < 0.0001, S4C Fig.). Both temperature (ANOVA: P < 0.0001) and pH (P < 0.0001) differed among soil samples by biome (S4B Fig., S4D Fig.). Soils from both hot and cold deserts tended to be basic while agricultural fields, forests, and tundra were acidic. Tundra and cold deserts were the coldest biomes, and shrubland, agriculture, and hot deserts were among the hottest biomes.

### Beta Diversity

Sediment, biofilm/mat, and inland water habitats displayed the highest beta diversity among geographic locations (geoclusters), whereas soil, air, and marine water exhibited the lowest beta diversity (PERMDISP: P < 0.0001, Fig. 1C). To test that these patterns were not influenced by unequal sampling (number of geoclusters) of the habitats, we subsampled the habitats (to 20 geoclusters per habitat) and retested the patterns. All 100 subsamplings produced the same beta-diversity rankings, and all models were significant, indicating that unequal sampling did not influence within-habitat beta-diversity. Within the soil habitat, beta-diversity did not differ by biome (P = 0.526, Fig. 1D).

Overall, bacterial community composition differed significantly by habitat (PERMANOVA: P = 0.001, Fig. 1E) and by biome for soils (P = 0.001, Fig. 1F). Because salinity influenced the community composition for both sediment (P = 0.002) and inland water (P = 0.002), we tested whether salinity likewise influenced beta diversity within sediments and inland water. Beta diversity did not differ between saline and non-saline samples within sediments (PERMDISP: P = 0.21) or inland water (P = 0.849, S5 Fig.).

The above results did not depend on the beta-diversity metric used. On a sample by sample basis, nine beta-diversity indices, including Bray-Curtis, were correlated with each other (r = 0.435 – 1.00, P = 0.001, mean r = 0.88, S6A Fig.). Raup-Crick has been suggested as a more appropriate metric when comparing groups with different alpha-diversity [33]. Because Raup-Crick assumes that all communities are part of the same regional species pool, we calculated the mean and standard error within each habitat (excluding between habitat comparisons) and compared the trend to that generated by the Bray-Curtis metric. Both Bray-Curtis and Raup-Crick metrics showed the same trend of beta-diversity among habitats (S6B Fig.).

### Gamma Diversity

Considering the accumulation of taxon richness across geoclusters, sediments exhibited the highest global gamma-diversity of any habitat, followed by soils and inland water (Fig. 4). The sediment rarefaction curve showed little sign of flattening out, indicating that most taxa are yet to be sampled. In contrast, the soil curve noticeably leveled off, even at a similar level of sampling. The gamma diversity of marine water, biofilms/mats, and air were not statistically distinguishable from one another but, as a group, exhibited lower global diversity than inland water, soils, and sediments.

**Fig 4.**
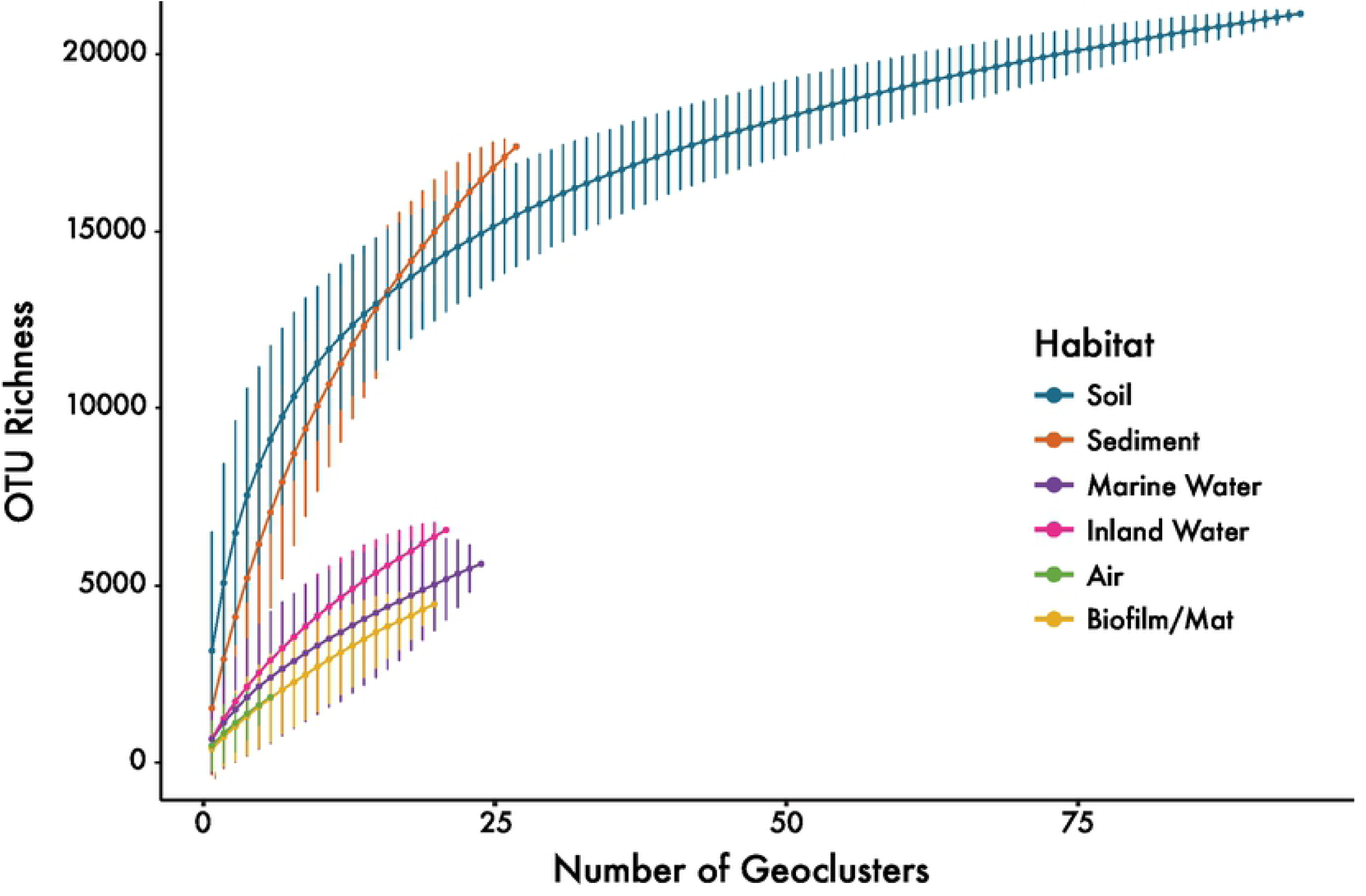
Geocluster accumulation curves. Geocluster accumulation curves (gamma-diversity) for mean OTU richness from a random sampling of geoclusters (permutations = 999) with 95% confidence intervals drawn for each habitat.

## Discussion

Here, we tested which habitat contains the most bacterial taxa within a single sample (alpha-diversity), which exhibits the most spatial variation among samples (beta-diversity), and which contains the most taxa on a global scale (gamma-diversity). We show that a single sample of soil on average contained higher bacterial alpha-diversity than any other habitat, including sediment (Fig. 1A). However, sediment contained more taxa globally, and much of its diversity is yet to be sampled (Fig. 6). Within soils, we found that agricultural soils had among the highest richness and exhibited just as much compositional variation (beta-diversity) as other biomes.

Although both sediment and soil were previously known to be highly diverse microbial habitats, previous studies demonstrated conflicting results about their relative ranking [14–17]. Using 11,680 samples and minimizing geographic biases, this analysis suggests that soil contains higher alpha-diversity than sediment. In contrast, marine water, inland water, air, and biofilms/mats contain the lowest alpha-diversity (Fig. 1A).

Within-habitat heterogeneity is a known driver of plant and animal diversity [34,35] and has been correlated with microbial communities as well [36,37]. Here, we show that the alpha-diversity patterns are consistent with the idea that habitat heterogeneity may drive bacterial diversity on a global scale. The highly mixed water and air environments harbor lower diversity, consistent with previous smaller-scale studies [16,38,39]. While both sediments and soil are not as well mixed, sediments contain higher water content than soils. Water content increases connectivity and thus reduces environmental heterogeneity and promotes dispersal, both of which can result in lower diversity [9,40,41]. At the same time, biofilms and mats, despite being spatially structured, also displayed low alpha-diversity [42]. However, the biofilm/mat samples from the EMP dataset encompassed samples with the highest pH and temperature (S3 Fig.). We therefore speculate that these abiotic extremes contribute to low alpha-diversity [43–45]. Yet ultimately, little is known about the environmental conditions, and their heterogeneity, at the spatial scale that matters for microorganisms [46]. To test the importance of within-sample heterogeneity on microbial diversity directly, finer-scale data are needed.

Our analysis is the first to quantify bacterial beta-diversity among habitats at a global scale. While soils contained the highest alpha-diversity within a single sample, sediments displayed higher beta-diversity among geoclusters samples habitat (Fig. 1C). Sediment beta-diversity was also similar to that of inland water and biofilms/mats. Additional environmental data associated with the individual samples would be needed to distinguish whether these beta-diversity patterns might be driven by dispersal limitation [9,47] or spatial variation in environmental conditions [6,38].

Given their high alpha- and beta-diversity, it is not surprising that sediments are also estimated to contain the highest global (gamma-) diversity (Fig. 4). The taxa accumulation curves also reveal that, while we may have observed most bacterial taxa in soil, we still have much more diversity to discover in sediments, at least with current sampling trends and methods.

Because plant diversity and productivity are shown to impact microbial communities [20,21], we further characterized alpha- and beta-diversity trends among biomes from which the soil samples were collected. Agricultural soils contained among the highest alpha-diversity, as previously noted in smaller scale studies [24,48]. Indeed, some agricultural practices, such as application of manure are known to increase bacterial diversity [49,50]. Even more notable, however, is that agricultural soils encompassed similar levels of beta-diversity to those of other biomes. While some agricultural practices have been shown to homogenize communities within a single field [48], not all practices have a homogenizing effect [51]. Further, the diversity of agricultural practices around the world [49,52] seems to select for as much variation in bacterial composition as forests or deserts.

Contrary to our hypothesis, biomes with higher plant diversity or productivity, such as forest or shrubland soils, were no more diverse within a sample than other biomes (Fig. 1B). These results support previous findings that grasslands contain more bacterial diversity than forests [53] and, overall, plant and soil diversity are uncoupled [54]. We therefore propose that abiotic factors may be more important for soil bacterial alpha-diversity than the biome. Biomes differed significantly in pH and temperature, and soil alpha-diversity was strongly correlated with both factors (S4 Fig.). These results are consistent with previous studies [12,55,56] that find bacterial richness in soils peaks at a neutral pH and at mid-temperatures. Of course, other environmental factors are likely influencing soil diversity. Soil water content, which increases connectivity among aqueous bacterial habitats, influences bacterial alpha-diversity as well, where intermediate soil water content promotes high diversity [41].

Finally, these diversity patterns appear to be robust to two key methodological issues. First, diversity trends did not depend on the particular alpha- or beta-diversity metrics used (Fig. 2A, S6A Fig.). Air, as the only exception, was more even than expected, given its richness (Fig. 2B). We speculate that the movement of air contributes to its evenness as air likely picks up a sampling of bacteria from many different habitats [57,58]. Second, the results were robust to the degree of clustering of the amplicon sequences (Fig. 3). Both the 97% OTU and the ESV datasets yielded the same alpha-diversity trends, as previously noted in a smaller scale study [59]. Most of those outliers in this analysis came from non-saline sediment or inland water samples, but we caution that these samples originated from only four geoclusters. Thus, while ESVs can be useful for resolving the diversity among specific groups [60], broad-scale alpha-diversity patterns do not seem to be altered by these particular operational definitions.

With the largest dataset created with consistent methodology and geographically widespread sampling effort, we show that soils support the highest diversity within a single sample (alpha-diversity) and that sediments are more variable in composition among locations (beta-diversity) and likely support the most bacterial taxa globally (gamma-diversity). Within soils, we find biome type impacts soil alpha-diversity but not beta-diversity. Many of these results appear consistent with the idea that spatial heterogeneity and dispersal limitation maintain bacterial diversity. These baseline patterns set the stage for new research on the mechanisms driving the generation and maintenance of bacterial diversity.

## Acknowledgements

We would like to thank Luke Thompson, Jack Gilbert, and the Earth Microbiome Project team for their vision and hard work in assembling this dataset and making it widely available. We also thank Steve Allison and Cascade Sorte for their input on analyses and methods, and Michaeline Albright and Alex Chase for help with statistical analyses and computational methods. We further thank Alex Chase, Cynthia Rodriguez, Sarai Finks, Claudia Weihe, Joia Capocchi, Kazuo Isobe, and Pauline Nguyen for their guidance with earlier drafts.

## Supporting Information

**S1 Fig.**
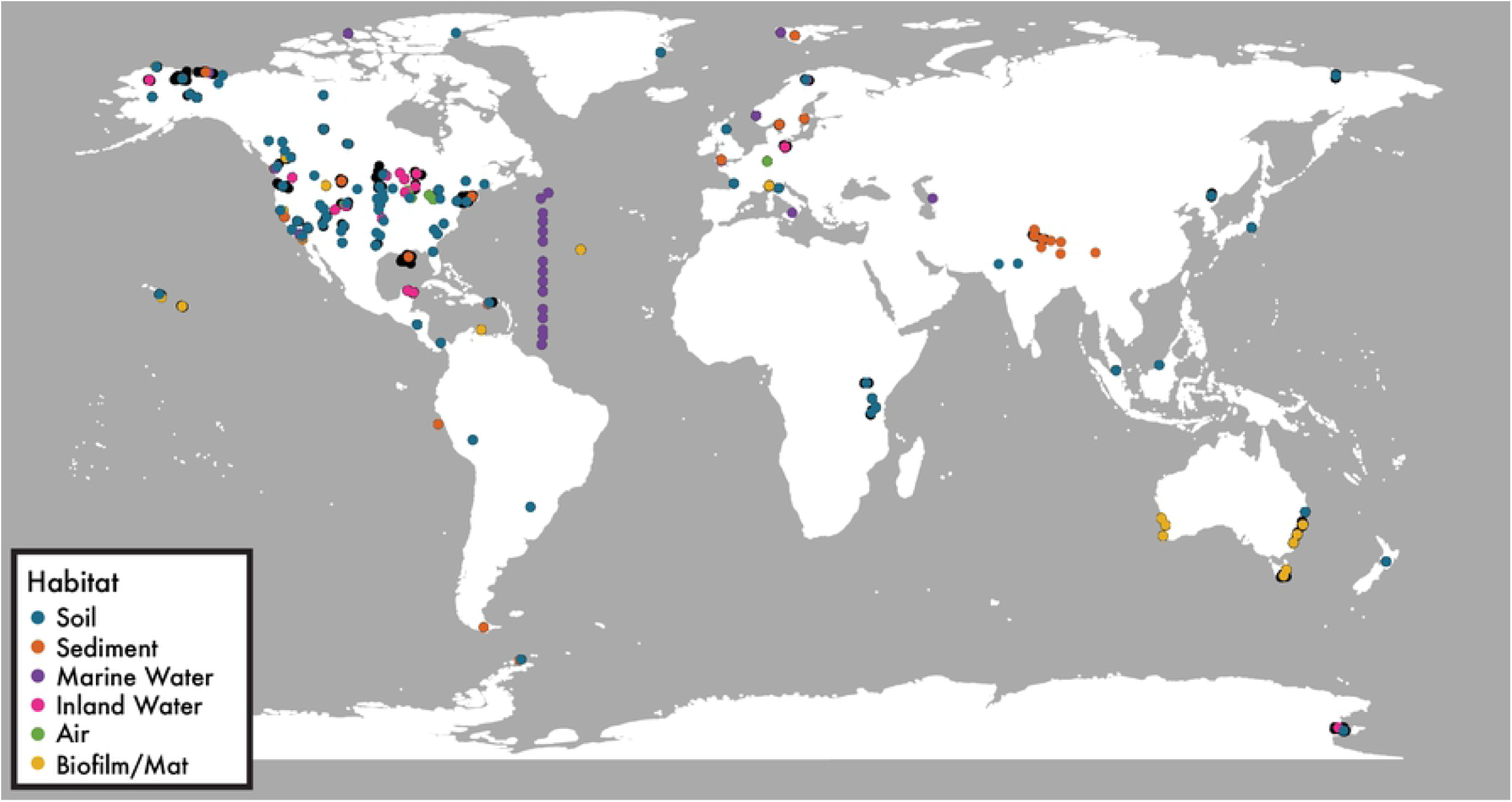
Sample and geoclusters locations. Map showing locations of each of the EMP samples used in this study (black dots) and the geoclustered samples for each habitat (colored dots). Geoclusters were created from samples located within 110 km of each other.

**S2 Fig.**
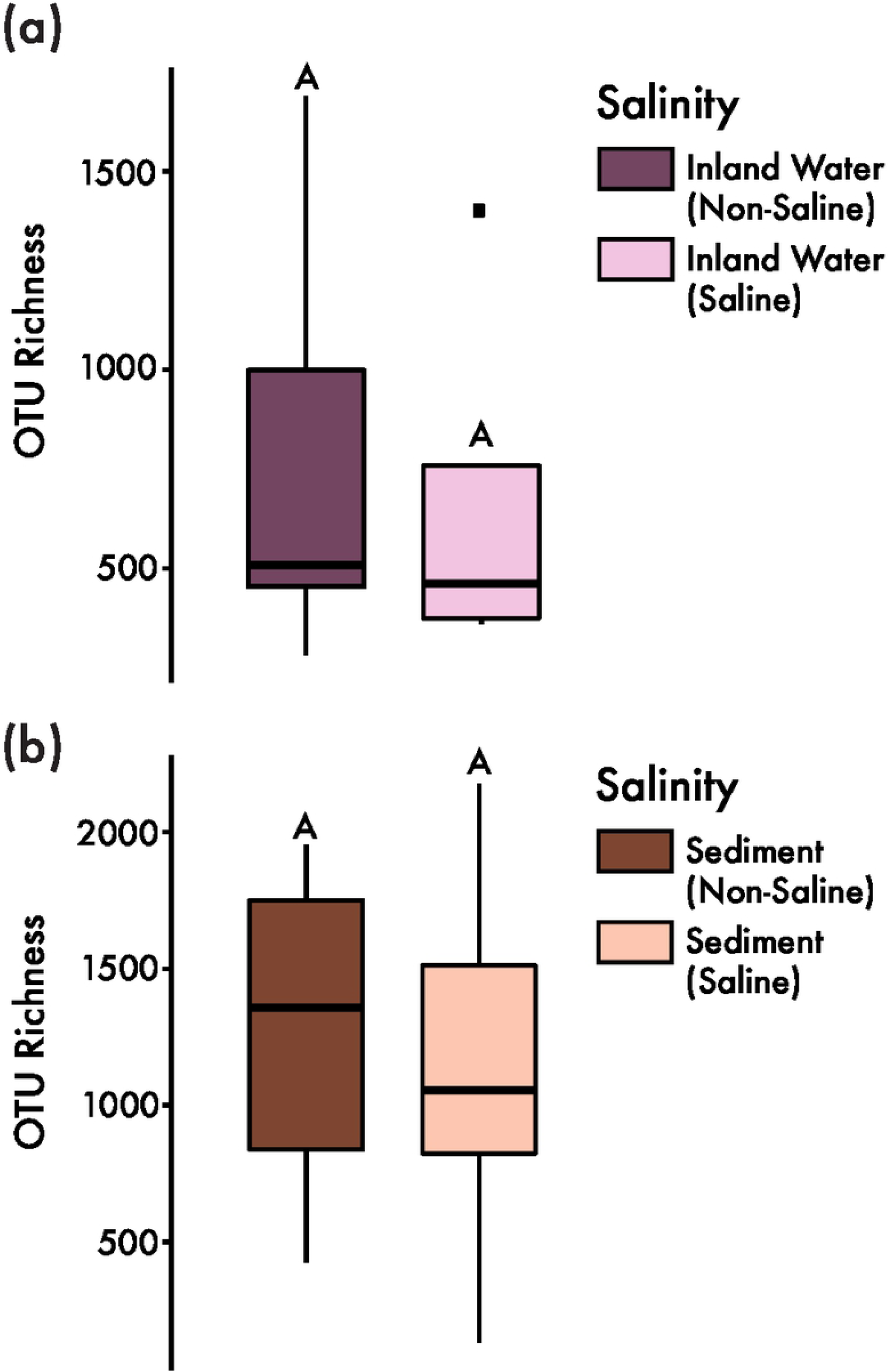
The influence of salinity on alpha-diversity. Taxon richness did not differ between saline and non-saline samples from inland water or sediment habitats.

**S3 Fig.**
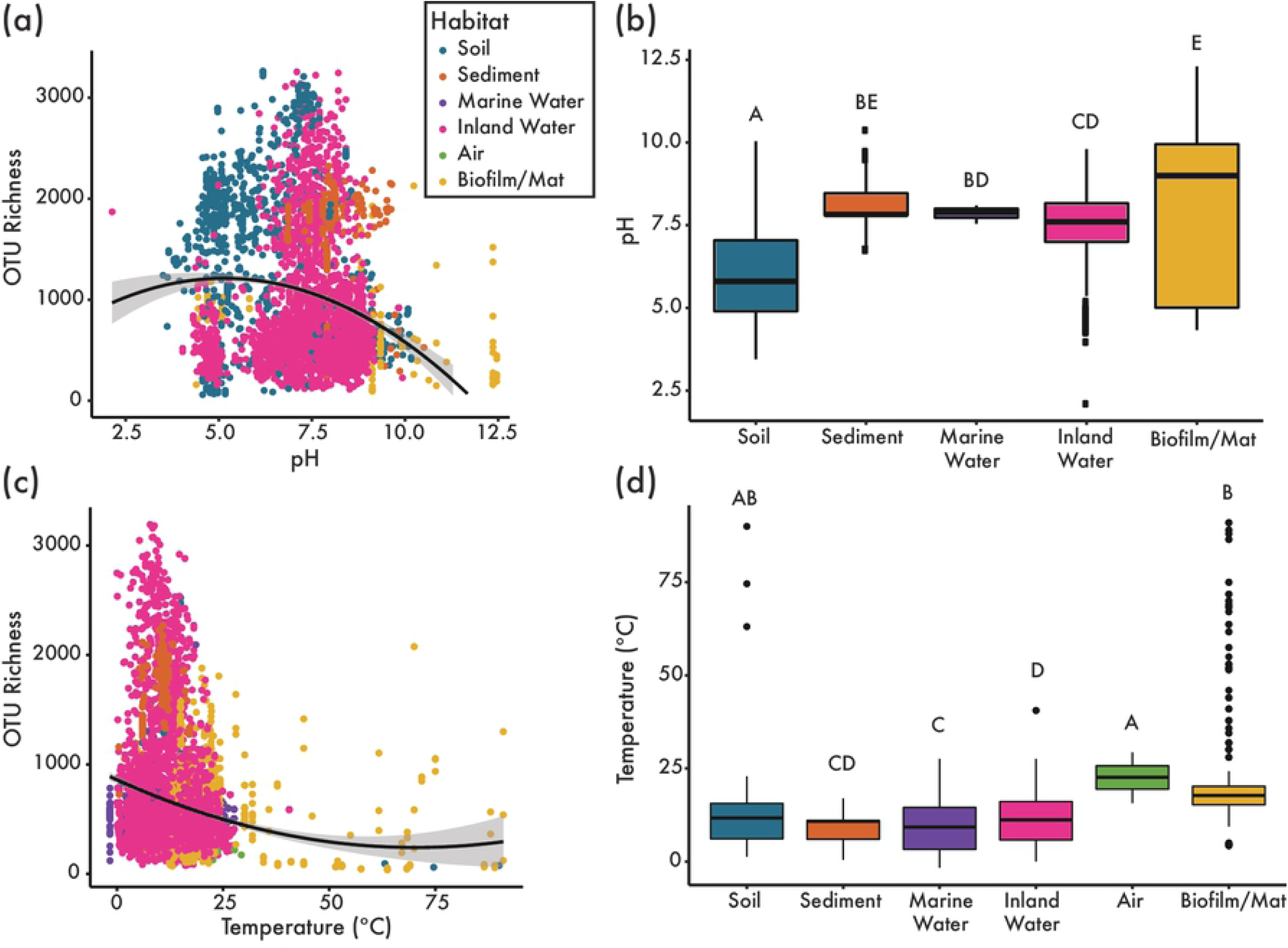
The influence of abiotic factors on taxon richness. (A) pH significantly impacts taxon richness (P < 0.0001). Each point represents an individual EMP sample (not a geoclusters) and is colored by habitat. (B) pH differs among habitats (ANOVA, p < 0.0001). Out of all the EMP environmental samples used in this study, 30.7% had associated pH metadata: 0% of air samples, 60.0% of inland water samples, 19.6% sediment samples, 6.5% of marine water samples, 5.1% of biofilm/mat samples, and 23.4% of soil samples. (C) Temperature significantly influences taxon richness (P < 0.0001). Each point represents an individual EMP sample (not a geoclusters) and is colored by habitat. (D) Temperature recorded when samples were collected (EMP metadata) differs among habitats (ANOVA, p < 0.0001). Out of all the EMP environmental samples used in this study, 47.3% had associated temperature metadata: 9.0% of air samples, 77.0% of inland water samples, 17.5% sediment samples, 48.8% of marine water samples, 81.0% of biofilm/mat samples, and 0.6% of soil samples.

**S4 Fig.**
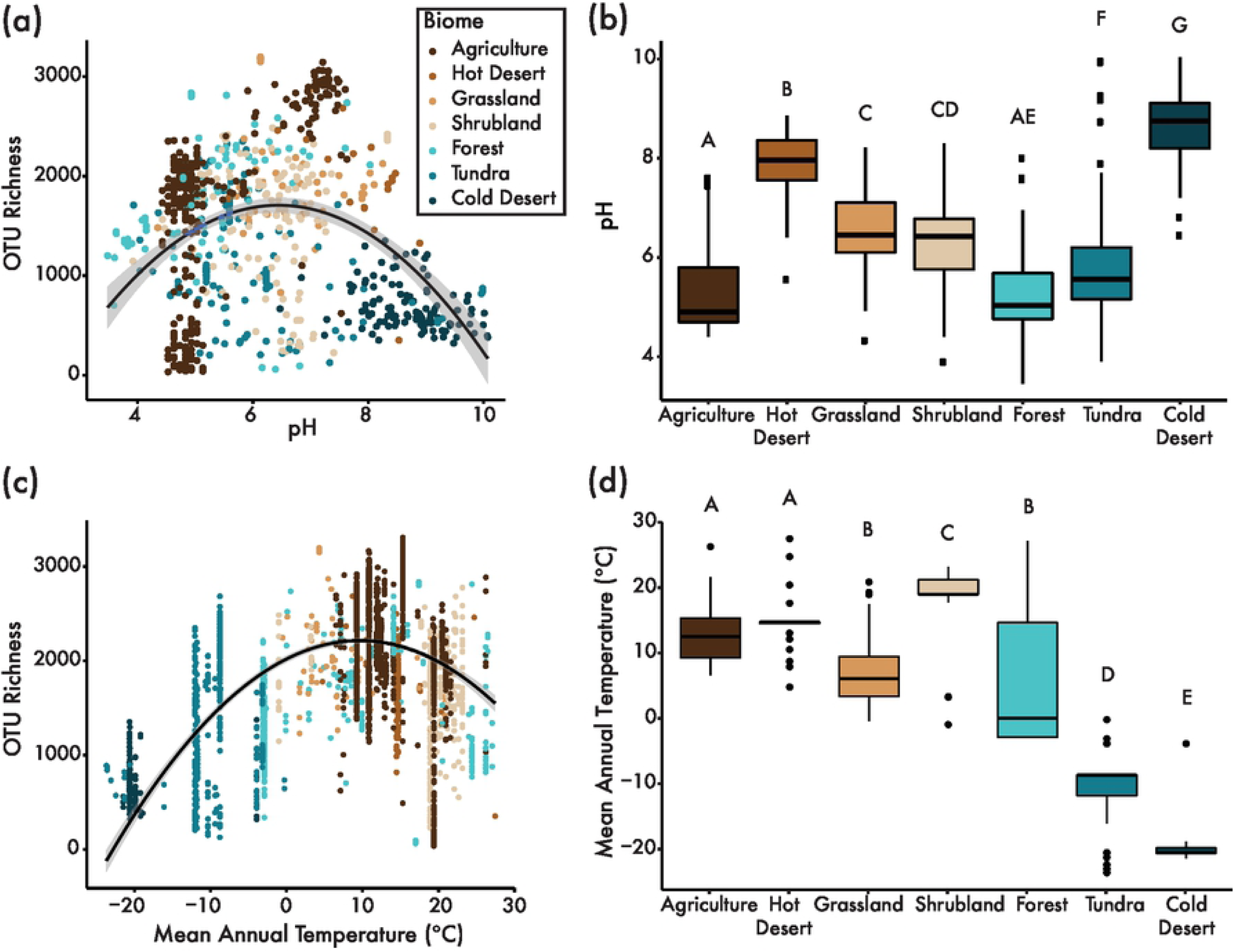
Abiotic factors influence taxon richness in soil. (A) pH significantly impacts taxon richness (P = 0.001). Each point represents an individual EMP soil sample (not a geocluster) and is colored by biome. (B) pH differs among biomes (ANOVA, p < 0.0001). (C) Mean annual temperature significantly impacts taxon richness (P < 0.0001). Each point represents an individual EMP soil sample (not a geocluster) and is colored by biome. Temperature data was retrieved from WorldClim, a publicly available data source. (D) Temperature differs among biomes (ANOVA, p < 0.0001).

**S5 Fig.**
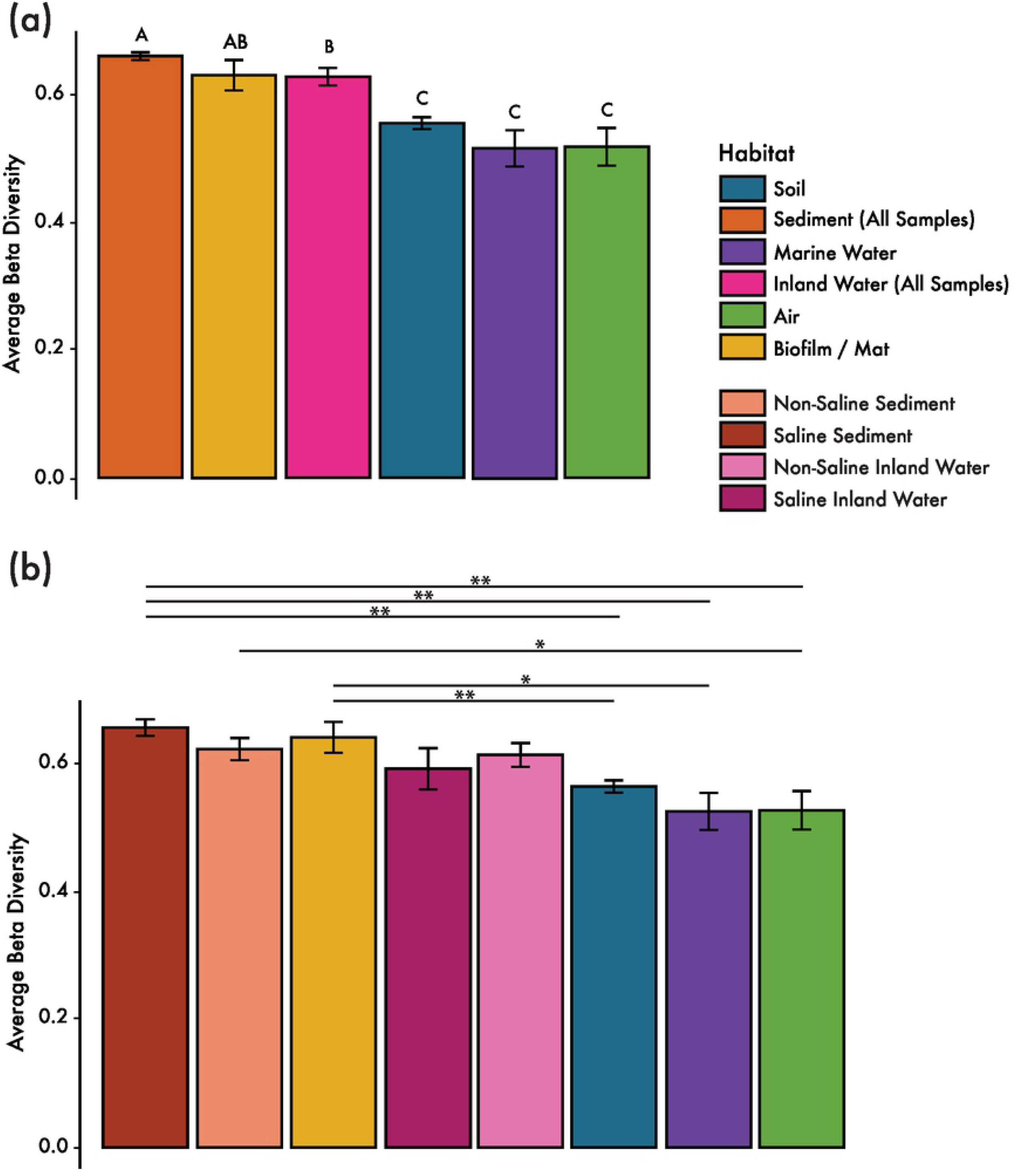
The influence of salinity on beta-diversity. The level of beta-diversity within sediment and inland water is not driven by the combination of saline and non-saline samples within a single habitat. (A) Mean beta-diversity (distance from centroid) ± standard deviations of the six habitats, ranking habitats from highest to lowest beta-diversity. (B) Mean beta-diversity ± standard deviations of all six habitats with sediment and inland water habitats split into saline and non-saline samples.

**S6 Fig.**
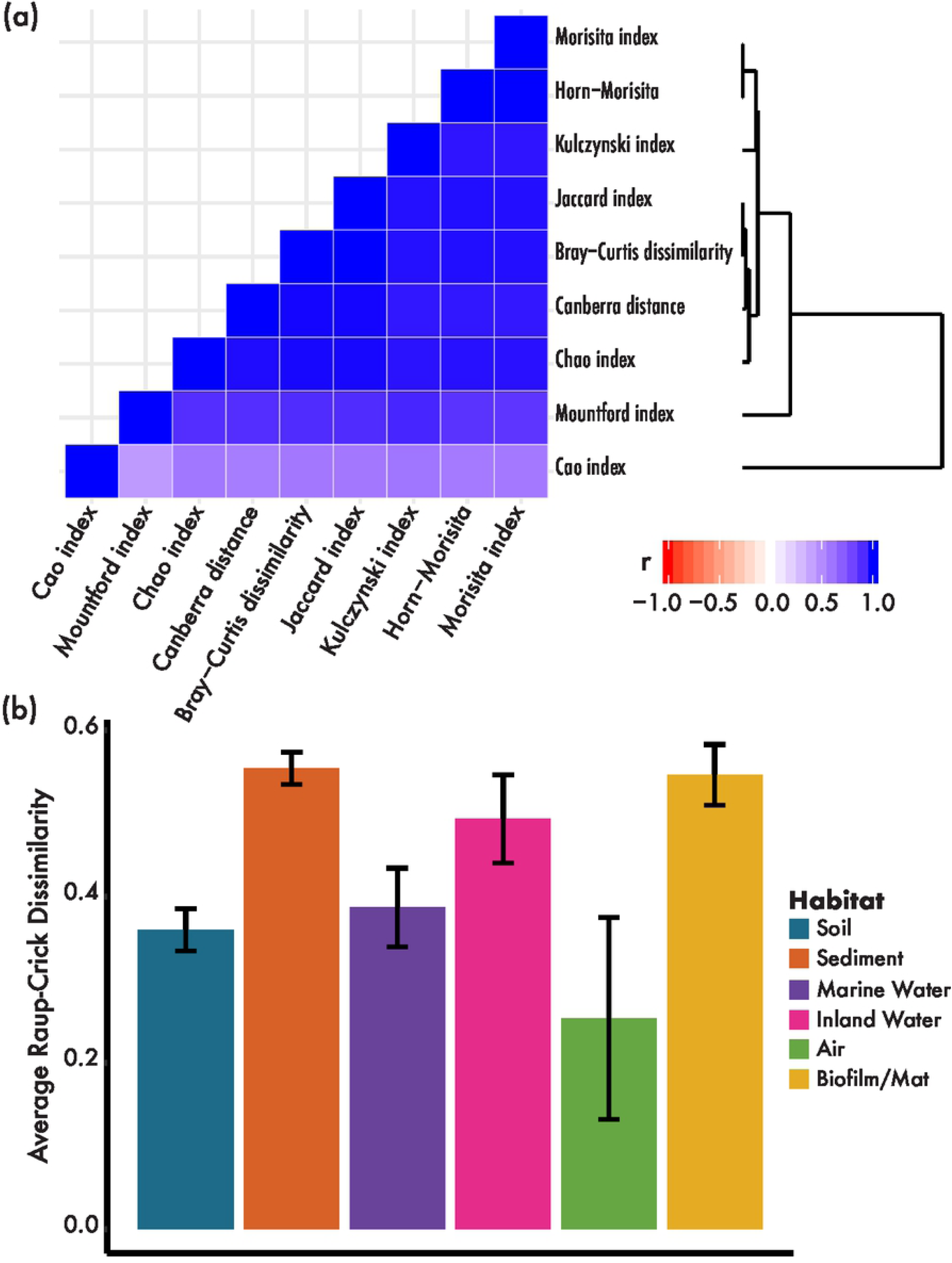
Comparison of beta-diversity metrics. Beta-diversity patterns are not dependent on the beta-diversity metric used. (A) Heatmap shows degree of correlation (r from a Spearman’s mantel test with all EMP samples used in analysis). Dendrogram shows relatedness of metrics based on their correlation strength. (B) Mean Raup-Crick dissimilarity (distance from centroid) ± standard deviations. The patterns shown by Raup-Crick analysis match those demonstrated by Bray-Curtis metric.

